# Structural determination of the HIV-1 Variable Region 3 epitope of antibody 19b

**DOI:** 10.64898/2025.12.17.694829

**Authors:** Susan K Fetics, Ariha Mehta, Nathan I. Nicely, Chun-Hsing (Josh) Chen, Jared Lindenberger, Priyamvada Acharya

## Abstract

The HIV-1 Envelope (Env) in its pre-receptor “closed” conformation is targeted by broadly neutralizing antibodies (bnAbs), while its receptor-bound “open” conformation, exposes immunodominant epitopes targeted by non-neutralizing antibodies. A human immunoglobin G (IgG) monoclonal antibody (mAb), 19b, binds an Env third variable (V3) loop epitope that is only exposed in the open Env conformation. Despite widespread use of 19b to detect the open Env conformation in immunoassays, its epitope has not yet been structurally defined. Here we determine crystal structures of ligand-free and V3 peptide-bound 19b Fab to visualize details of this interaction. 19b utilizes both its heavy and light chains to interact with the V3 loop. The 5-residue heavy chain complementarity-determining region (CDR H3) facilitates a hydrophobic pocket for V3 residues to associate with. 19b adopts a cradle binding mode with its CDRH1, CDRL2 and CDRL3 mediating interactions with V3 regions that flank the conserved GPGR/Q motif, without making substantial contacts with the GPGR arch region. Our high-resolution structures by elucidating the epitope, binding mode and the structural basis for the broad reactivity of 19b, fill a gap in our knowledge of a reagent that is widely used in immunoassays.

## Introduction

Human immunodeficiency virus type 1 (HIV-1) expresses a single viral protein, called envelope (Env), on its cell surface. As a result, Env is the sole target for HIV-1 neutralizing antibodies (1). The Env glycoprotein has five variable loop regions (V1–V5), of which the V3 loop harbors conserved binding motifs at its base and a GPGR/Q motif at its apex that interact with the coreceptor, CCR5 or CXCR4. The sequence of V3 predicts co-receptor specificity while its conformation determines co-receptor binding and virus infection (2, 3).

In the “closed” Env conformation prior to receptor binding the V3, shielded by its interactions with the V1V2 regions, is inaccessible for co-receptor binding. After Env engages the CD4 receptor, conformational changes occur in Env that are collectively termed as Env “opening”. In the open Env conformation, the V3 region extends out from the Env core, no longer interacting with V1V2, with the extended V3 available for coreceptor binding. Antibodies that target the V3 crown in the open Env conformation are non-neutralizing or show strain-specific neutralization and are present in most persons infected with HIV-1 (**Figure 1A**) (4).

**Figure 1.**
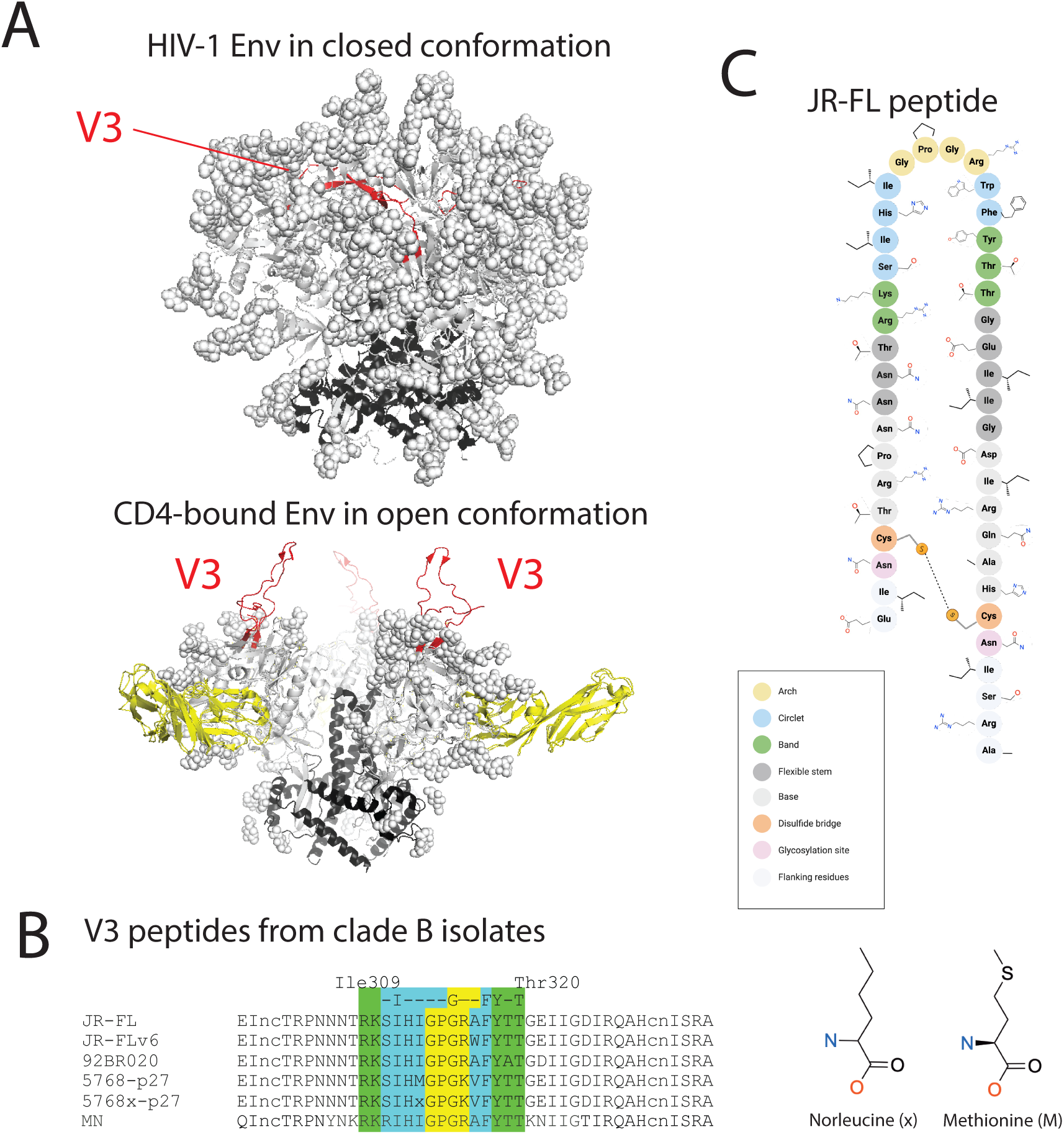
Crystal structure of 19b Fab in the presence of peptide V3 JR-FL. (A) HIV-1 Env shown in cartoon representation in the closed (top) and CD4-bound open (bottom) conformations, with gp41 colored black, gp120 grey and the V3 peptide colored red. Glycans are shown as spheres. (B) V3 peptide sequences used in this study and of isolate MN. (C) Key side chain interactions between 19b Fab and V3 JR-FL residues Arg306, Arg315, Thr319 and Thr320. (D) Electrostatic potential surface of (left) V3 JR-FL peptide and (right) 19b Fab epitope. Blue represents positive charge; red denotes negative charge.

The V3 region is ∼50 Å in length and consists of a conserved base (residues 296-300 & 326-331), a flexible stem (residues 301-305 & 321-325) and a β-hairpin loop crown (residues 306-320, **Figure 1B**). A disulfide bond between residues C296 and C331 at the base stabilizes the V3 region (2, 5, 6) (**Figure 1C**). The crown can be further broken down into three regions: the band (306 – 308 & 318 - 320), the circlet (309 – 311 & 316 – 317) and the arch (GPGR/Q at 312 – 315), with the band and circlet regions flanking the GPGR/Q arch (7).

Human IgG monoclonal antibody N701.9b, commonly known as 19b, was isolated from an asymptomatic person with HIV-1 (8, 9). Alanine scanning of V3 residues 308 to 318 showed that Env binding to 19b occurs at residues 313 – 318 within the crown (8). Further amino acid substitution analysis revealed a V3 motif that directly affects Env interaction with 19b: -I G--FY-T, where dash denotes residue of any identity and letter denotes the residue where a contact must occur. These residues span the V3 β-hairpin turn, occupying the arch, circlet, and band regions. Despite the diversity of V3 residues in HIV-1 strains, 19b binding is observed for HIV-1 subtypes A, B, C, E and F, illustrating inter-clade cross-reactivity (9, 10). As a poorly neutralizing antibody, 19b binds only neutralizes Tier 1 isolates (11). Early studies indicated that the Env glycan shield can prevent accessibility of the 19b epitope (12).

Here, to define the 19b epitope at the atomic level, we deteremined crystal structures of the 19b antigen binding fragment (Fab) in complex with aglycone V3 peptides from HIV-1 clade B isolates JR-FL, 92BR020 and 5768-p27. 19b neutralized the 5768-p27 and 92BR020 isolates with IC_50_ of 3.2 ug/ml and 21.5 ug/ml, respectively, while no neutralization of JR-FL was observed at the concentrations tested (IC_50_ >50 ug/ml) (13). We also determined a structure of the ligand-free 19b Fab. 19b engages the V3 in the cradle binding mode (abbreviated as V3C) with contact made with the hydrophobic core residues in the circlet and the band regions, with minimal contacts with the GPGR/Q arch. The cradle binding mode adopted by 19b is distinguished from the ladle binding mode (V3L) where interactions occur with the GPGR arch. The 5-residue complimentary-determining region (CDR) 3 in the heavy chain (CDR H3) of 19b creates a pocket for V3 binding. The dominant interactions are established between the band and circlet regions of V3 and the CDR 2 and 3 of the light chain (L2 and L3) and the CDR H1.

## Results

### Crystallization of 19b Fab with HIV-1 Env V3 peptides

For co-crystallizing with 19b Fab, we synthesized V3 peptides from clade B HIV-1 isolates JR-FL, 92BR020 and 5768-p27 (**Figure 1B**). A modified version of the JR-FL V3 peptide, called JR-FL.v6 V3 peptide here, was used (14). In addition to the native 5768-p27 V3 peptide, a second peptide called 5768x-p27 was synthesized, where Met311 was substituted with norleucine to prevent unwanted oxidation (**Figure 1C**). The synthetic peptides spanned 43 residues (295–337), including the V3 base, crown and stem regions, with a disulfide bond between Cys297-Cys332.

19b Fab was crystallized, ligand-free and bound to four aglycone V3 peptides. Each crystal data set contains 1 molecule in the asymmetric unit with the space group P2_1_2_1_2_1_ and similar unit cell parameters (**Table S1, S2** and **Figure 2**). For this study, residue numbering for 19b follows the Kabat and Wu convention and for the V3 peptides follows the HXB2 number scheme (15). In the 19b-bound V3 peptide structures electron density was present for the central 15 – 19 residues of V3 flanking the GPGR/K arch, wherein the peptide model was built (**Figure 2A**). All the bound V3 peptides showed similar β-hairpin loop conformation, exhibiting a type-II β turn at the GPGR/K arch residues 312-315, stabilized by hydrogen bonds mediated by backbone residues (**Figure 2B**). Superposition of the V3 peptides from the different structures yielded Cα RMSD values of ∼0.2 Å (**Table S3**).

**Figure 2.**
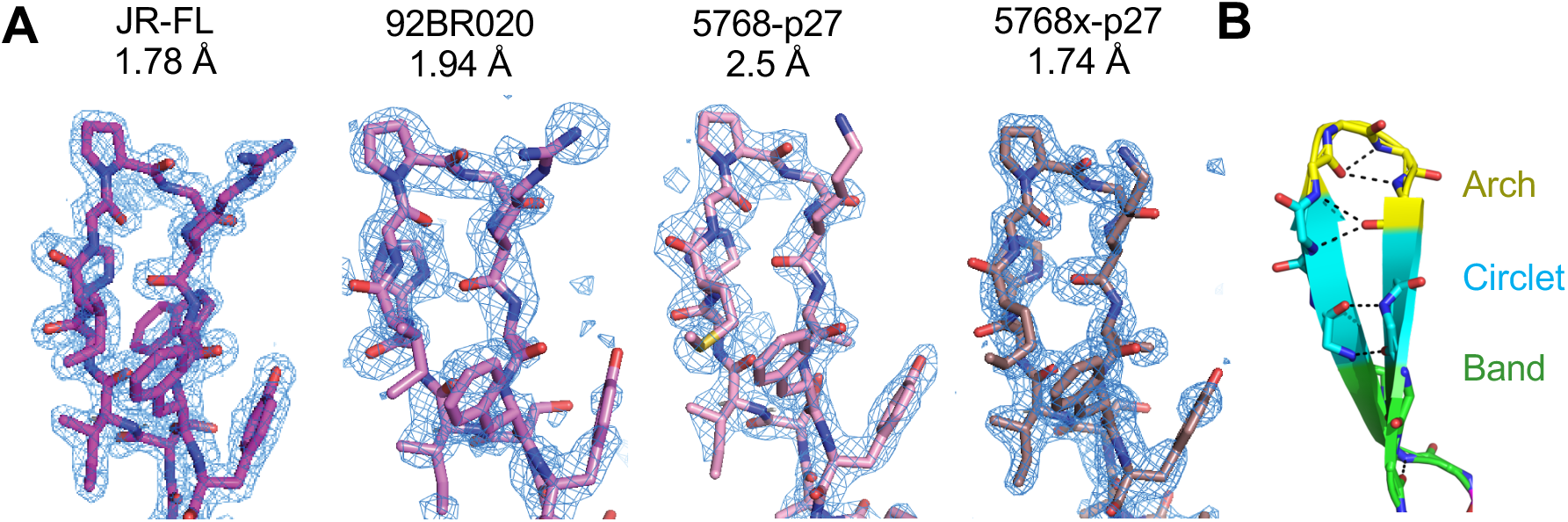
Crystal structure of diverse clade B V3 peptides bound to antibody 19b. (A) Difference density represented as blue mesh of V3 peptides JR-FL (magenta), 92BR020 (violet) 5768-p27 (pink), 5768x-p27 (dark violet). Fo-Fc map is contoured at 3α. (B) The 19b-bound JR-FL peptide main chain is shown in cartoon representation and colored by the arch (yellow), circlet (cyan) and band (green) regions, and and representation colored by atom. Dashed lines indicate hydrogen bonds.

### Details of V3 peptide interactions with 19b

To visualize the details of the interactions of 19b with the HIV-1 Env V3 region, we first studied the JR-FL.v6 V3 peptide-bound 19b Fab complex (**Figure 3**). Both heavy and light chains of 19b were involved in interactions with V3, with the 19b CDRs contacting the V3 crown at the interface of the antibody heavy and light chains (**Figure 3A and 3B**). The 19b light chain contributed a more electronegative surface compared to the heavy chain (**Figure 3B**). Electrostatic complementarity was observed in the interaction as the more electropositive surface of the V3 peptide interacted with the electronegative region of the 19b paratope (**Figure 3B**). The V3 crown interacted with four of six 19b CDRs: L2, L3, H1 and H3 (**Figure 3C-E**). All three regions of the V3 crown – the arch, circlet, and band – contacted 19b (**Figure 3C-E**). Characteristic of the cradle binding motif, the GPG turn of the GPGR arch motif was solvent exposed (**Figure S1**). While this region did not make direct contacts with 19b, hydrogen bonds mediated by a solvent network were observed with heavy chain residues Asp31 (H1), Tyr32 (H1) and Val100 (H3), and light chain residues Ser52 and Glu55 (L2) (**Figure 3C**). A salt bridge was observed between the side chain guanidine group of Arg315 of the GPGR motif with the 19b residue Glu55 (L2). The 19b CDR L2 residue Glu55 contributes to the electronegative character of the portion of the paratope coming from the 19b light chain (**Figure 3B**). The side chain of Arg315 was further stabilized by stacking against the side chain of Tyr49 (L2) (**Figure 3C**).

**Figure 3.**
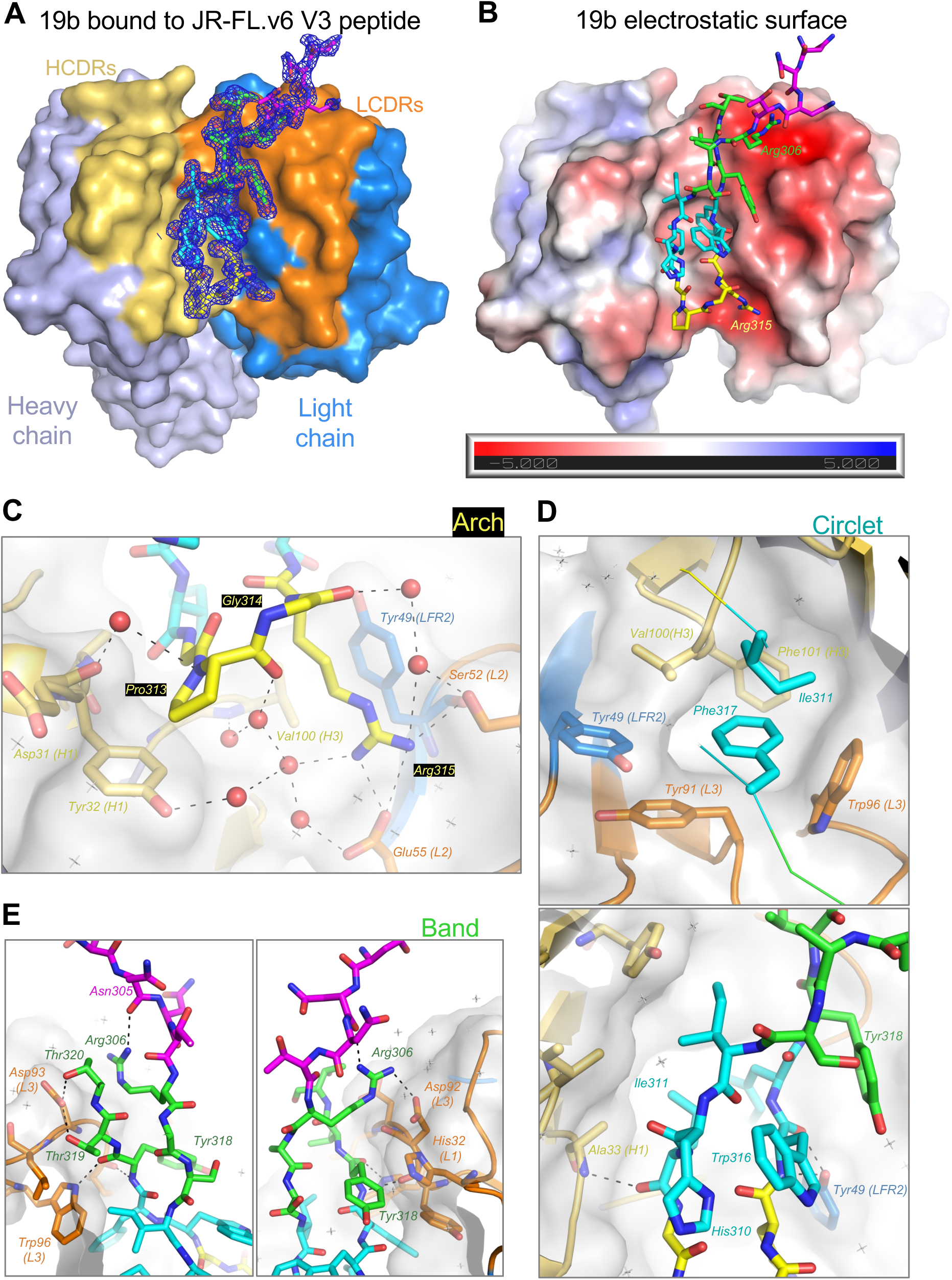
Crystal structure of 19b Fab in complex with the JR-FL.v6 V3 peptide. (A) 19b Fab shown in surface representation bound to the JR-FL V3 peptide shown in stick representation. 19b Fab heavy chain is colored light blue and light chain is marine. The CDR loops are colored: L1/L2/(L3 (orange), H1/H2/H3 (yellow-orange). The cradle motif of epitope interaction for V3 and 19b Fab is observed. Electron density for V3 peptide of JR-FL is shown as blue mesh with 2Fo-Fc map contoured at 1α. JR-FL V3 is shown in stick representation with atoms colored as follows: band (green), circlet (cyan), arch (yellow). Residues outside these three regions in the V3 loop are colored magenta. (B) Electrostatic potential surface of 19b Fab with bound V3 peptide shown as sticks and colored similarly as in panel A. (C-E) Details of interactions between 19b Fab with V3 residues in the (C) Arch, (D) Circlet and (E) Band regions. Two ∼180° rotated views to illustrate the interactions of the Band region with 19b. 19b is shown in cartoon representation with V3-interacting residues shown as sticks. An overlaid light gray transparent surface is also shown for 19b. Dashed lines indicate hydrogen bonds. Key water molecules are shown as red spheres. Other waters are visible as light grey “+” signs.

A key interaction was mediated by the V3 circlet residue Phe317 that inserts its aromatic side chain into a pocket lined by hydrophobic and aromatic residues – Tyr91 (L3), Trp6 (L3), Tyr49 (LFR2), Val 100 (H3), Phe101 (H3) and V3 residue Ile312 (**Figure 3D**). The central role of residue Phe317 in V3 binding to 19b was recognized previously (13). Other circlet interactions include hydrogen bonds mediated by the main chain of residues 311 and 316. The main chain carbonyl of Ile311 engages in a hydrogen bond with the main chain nitrogen of Ala33 (H1). The 19b light chain residue Tyr49 (LFR2) utilizes its side chain hydroxyl to engage in hydrogen bonds with the main chain of residue 316. Residue 316 in the JRFL.v6 V3 peptide is the site of the Ala to Trp substitution that was incorporated to stabilize the pre-fusion, closed Env conformation (14). The lack of interaction of the Trp316 side chain with the antibody explains why binding of the V3 peptide to 19b is compatible with the substitution of Ala316 in JR-FL Env to the bulkier Trp. In our structure, the Trp side chain stacks against the side chain of V3 residue His312.

The band segment of the V3 crown uses both main chain and side chain contacts to interact with 19b (**Figure 3E**). The main chain carbonyl of V3 residue Tyr318 hydrogen bonds with the 19b residue Trp96 (L3), while the Asp93 (L3) side chain engages in hydrogen bonds with the side chains of the V3 residues Thr319 and Thr320. An intra-V3 hydrogen bond was observed between the side chain of residue Arg306 and the main chain carbonyl oxygen of Asn305. This interaction brings the positively charged side chain of Arg306 in the proximity of the electronegative surface contributed by the light chain of 19b and facilitates a salt bridge interaction with Asp92 (L3) (**Figure 3B and 3E**).

In summary, the crystal structure of 19b Fab bound to the JRFL.v6 V3 peptide provides atomic level visualization of the V3-antibody interactions and provides structural basis for the previously observed roles of critical residues involved in the interaction.

### Recognition of diverse V3 peptides by 19b

To obtain quantitative measures for the affinity and kinetics of the interaction of 19b with the V3 peptides we performed surface plasmon resonance (SPR) binding assays (**Figure 4A and B**). The 19b Fab bound the JR-FL.v6 and 92BR020 V3 peptides with a similar affinity of 1.74 nM (**Figure 4B**). The V3 sequences of these isolates differ by two residues at positions 316 and 319 (**Figure 1B**). The 5767-p27 V3 peptides showed ∼20-fold tighter binding (K_D_ = 0.087 nM), with a substantially slower off-rate compared to the JR-FL.v6 and 92BR020 V3 peptides. Therefore, despite our crystal structures showing remarkable similarity in the 19b-bound conformations of V3 peptides from different HIV-1 isolates (**Figure 2A**), we observed considerable variation in the affinity and kinetics of binding of the V3 peptides to 19b. The differences in affinity observed between the 92BR020 and the 5767-p27 V3 peptides trended with the differences observed in the neutralization of the corresponding pseudoviruses by 19b (13). On the other hand, JR-FL was not neutralized by 19b even though its V3 peptide bound with similar affinity as the 92BR020 V3 peptide, consistent with the JR-FL isolate predominantly being in the closed conformation with the V3 crown epitope occluded and inaccessible for 19b binding.

**Figure 4.**
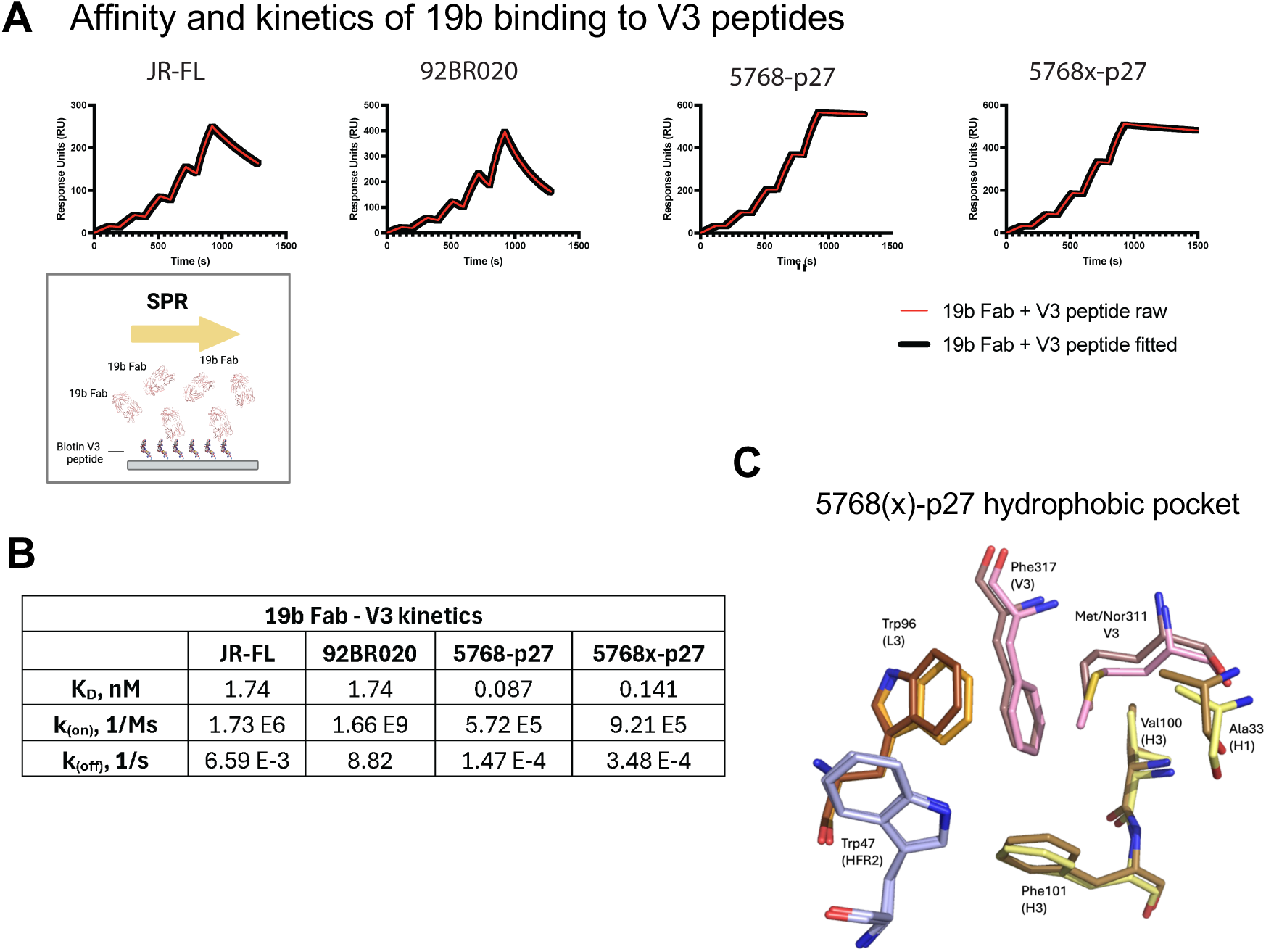
19b binding to diverse V3 peptides. (A) SPR sensorgrams shown for 19b Fab binding to V3 peptides from (left to right), JR-FL, 92BR020, 5768-p27, and the 5768x-p27 V3 peptide where the methionine at position 311 was replaced with a norleucine in the 5768-p27 peptide. The reference subtracted sensorgrams are shown as red lines in the graphs with the black lines indicating fit of the data to a 1:1 Langmuir binding model. The inset is a schematic of the experimental design showing the V3 peptide captured onto a streptavidin surface via its biotin tag and the 19b Fab flowed over the surface to measure binding. (B) Table listing the affinity and kinetics of binding of each V3 peptide to the 19b Fab. (C) Hydrophobic pocket that into which V3 Phe 317 gets buried in its 19b-bound state is shown for the 19b complexes with the 5768-p27 and 5768x-p27 V3 structures. For the 5768-p27 structure, V3 is colored pink, LCDR3 bright orange, HCDR1 and HCDR3 pale yellow, and for the 5768x-p27 structure, V3 is colored dark violet, LCDR3 brown, HCDR1 and HCDR3 sand.

In the 5768(x)-p27 – 19b Fab structures, the methionine (in 5768-p27) and the non-standard norleucine (in 5768x-p27) replaces Ile at position 311 to participate in forming the hydrophobic pocket that buries the V3 Phe317 residue (**Figure 4C**). The preference for a Met at V3 position 311 for its interaction with 19b was discussed previously (13). Indeed, this is reflected in the ∼10-fold enhanced affinity for the 5768-p27 V3 peptides for the 19b Fab compared to the JR-FL and 92BR020 V3 peptides. The V3 of 5768(x)-p27 harbors a lysine at position 315 within the arch in lieu of arginine in JR-FL and 92BR020, thereby weakening the interaction observed between V3 position 315 and Glu55 (L2) of 19b as seen in JR-FL and 92BR020 (**Figure 3C**), confirming the non-essential nature of this arch interaction and further supporting the circlet/band “cradle” binding motif between 19b Fab and Env V3 peptide (**Figure S1**).

### Crystal structure of ligand-free 19b Fab

To study V3 peptide binding induced changes in 19b, we solved the crystal structure of V3-free 19b and compared it to the structure of 19b bound to the JR-FL V3 peptide (**Figure 5**). We observed close agreement between the two structures with Cα RMSD value of 0.807 Å (**Figure 5A**). The six CDR main chains adopt the same conformation in the ligand-free 19b when compared to 19b in complex with V3 JR-FL. The Cα RMSD values for all six CDRs between the ligand-free 19b Fab and 19b Fab co-crystallized with V3 JR-FL structures are below 0.3 Å (**Table S3**). Ligand-free 19b CDRs L2, L3 and H1 have Cα RMSD values of 0.143 Å, 0.117 Å, and 0.069 Å respectively compared to the JR-FL V3 – 19b structure (**Table S3**). Around the 19b L2 region two water molecules were observed that formed hydrogen bonds with the main chain carbonyls of residues Ser52 and Val51 (**Figure S2**). Testing whether the protein structure prediction algorithm, AlphaFold can recapitulate the V3 peptide-antibody binding interface observed in the crystal structures, notable divergence in the main chain occurs at CDR L2 where the Cα RMSD is 1.456 Å between the experimental and Alfafold3 modeled structures, impacting the observed interactions of the V3 peptide with the L2 region (**Figure S2**). L2 Residue Ser52 in the Alfafold3 model was not within hydrogen bonding distance of Arg315 from V3 JR-FL, although the interaction between Glu55 (L2) and Arg315 (V3) was retained. These observations suggest a structural role for these waters in stabilizing the experimentally observed conformation of the L2 region.

**Figure 5.**
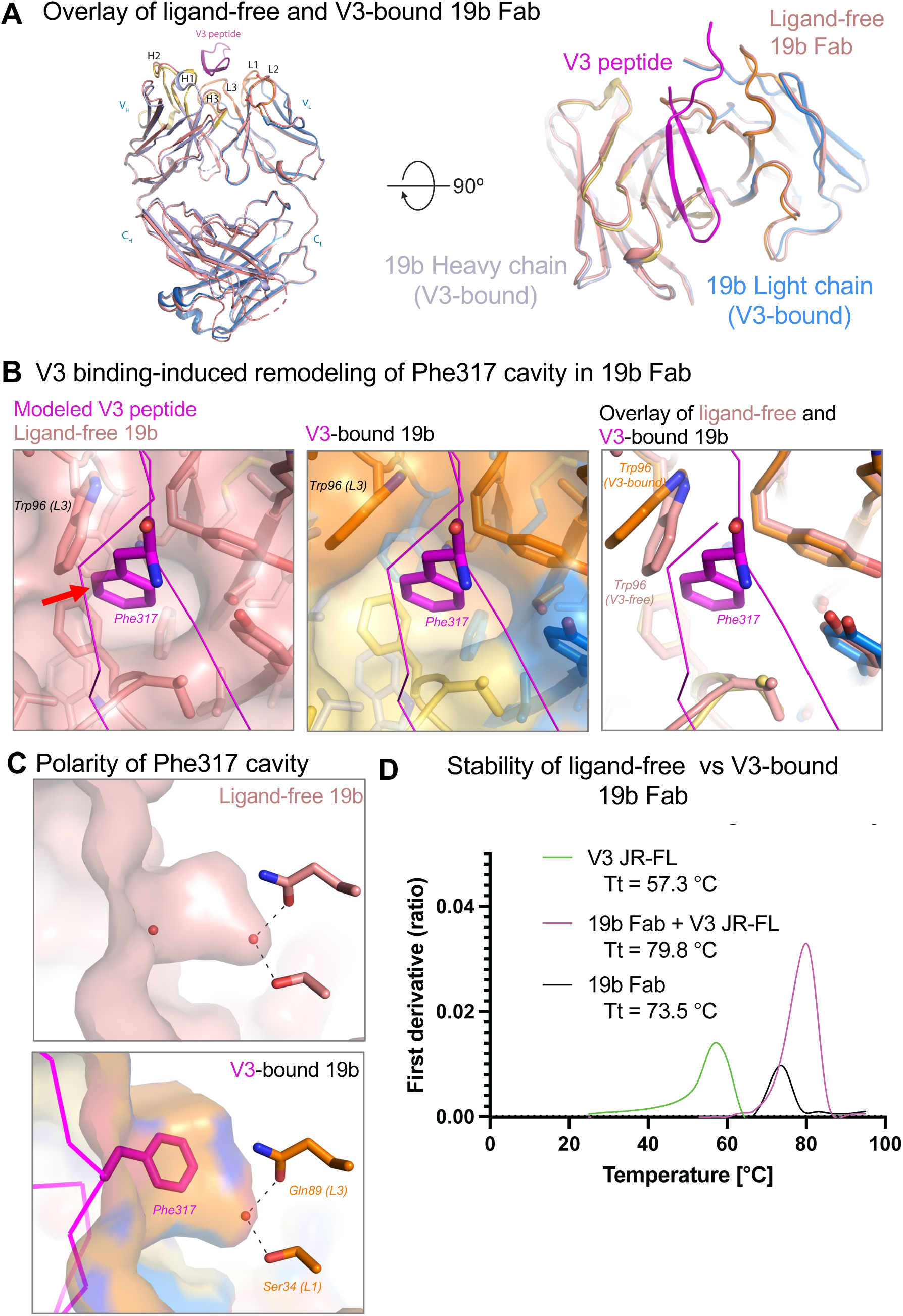
Crystal structure of ligand-free 19b Fab. (A) Crystal structure of the ligand-free 19b Fab (colored salmon) overlaid on the structure of the JR-FL.v6 V3 peptide (colored magenta) –bound 19b Fab (heavy chain colored light blue, light chain marine, HCDRs colored sand and LCDRs colored orange). Two 90° rotated views are shown. (B) Comparison of the cavity into which the V3 residue Phe317 side chain buries. Left. Ligand-free 19b structure shown as a transparent salmon surface and in cartoon representation with side chains shown as sticks. Shown in magenta ribbon representation is the V3 peptide with the Phe317 residue shown in stick representation. The V3 peptide is modeled onto the ligand-free structure, based on the overlay shown in panel A. The red arrow indicated a clash of the inserting Phe43 phenyl group with the wall of the cavity lined by the Trp96 (L3) side chain. Middle. JR-FL.v6 V3 peptide bound 19b, with the antibody structure shown as a transparent surface, and in cartoon representation with side chains shown as sticks. Right. Overlay of ligand-free and V3-bound structures zoomed into the Phe317 binding region. A rotameric shift in the Trp96 (L3) side chain enlarges the cavity allowing the phenyl group of the V3 residue Phe317 to insert. (C) Side views of the Phe317 insertion cavity. Top. Ligand-free 19b. Bottom. V3-bound 19b. Water molecules in the cavity are shown as red spheres. Dashed lines indicate hydrogen bonds. (D) Differential Scanning Fluorimetry of ligand-free 19b Fab (black), V3-bound 19b Fab (magenta) and of the V3 peptide (green).

Although the main chain configurations of the ligand-free 19b Fab and the V3-bound 19b Fab were similar, an important structural shift was observed in the “Phe317 cavity”, the binding pocket that is lined by 19b heavy and light chain residues, and into which the V3 residue Phe317 inserts (**Figure 5B**). In the V3 peptide-bound structure, a rotameric shift in the light chain Trp96 (L3) side chain enlarges the cavity allowing the phenyl side chain of the V3 Phe317 residue to insert. In the ligand-free 19b structure we observed two water molecules in this Phe317 insertion cavity – one near the mouth of the cavity and another placed deeper into the cavity making hydrogen bonds with light chain resides Ser34 (L1) and Gln89 (L3) (**Figure 5C**). In the V3-peptide bound structure, the water near the mouth of the cavity was displaced by the binding ligand but the second water molecule that was located deeper in the cavity was retained. Previous studies have shown that Phe317 can be substituted by Ile, Leu, Val and Tyr and still retain 19b binding (13). It is possible that when substituted by a Tyr, the Tyrosine side chain hydroxyl displaces the water from the cavity and takes its place to hydrogen bond with Ser34 (L1) and Gln89 (L3). Our structures thus reveal a malleable cavity that can adjust its shape to accommodate the size, shape and polarity of the incoming residue.

Using Differential Scanning Fluorimetry (DSF) to monitor changes in intrinsic protein fluorescence upon application of a thermal ramp, we observed an increase in the stability of the 19b Fab when it was bound to the V3 peptide compared to the ligand-free 19b Fab (**Figure 5D**). The ligand-free 19b Fab showed a transition temperature of 73.5 °C, which increased by 6.3 °C resulting in a transition temperature of 79.8 °C when the 19b Fab was bound to the JR-FL.v6 V3 peptide.

Taken together, these results show how the HIV-1 Env V3 engages with the 19b antibody and stabilizes it with minimal changes to the 19b main chain residues and elucidates the mechanism by which a key binding cavity for the V3 residue 317 adapts to V3 diversity and accommodates different amino acids.

## Discussion

The HIV-1 Env V3 region facilitates binding with the GPCR coreceptor, either CCR5 or CXCR4, which constitutes a key step in the entry of HIV-1 into host cells. Despite its variability across different HIV-1 isolates, key V3 loop motifs are conserved to enable its binding to coreceptor. The V3 region is targeted by mainly two categories of antibodies – the first that bind an epitope at the base of the V3 loop and the surrounding glycans, and the second that bind the V3 tip or crown region. 19b belongs to the second category of non-neutralizing or poorly neutralizing antibodies, whereas antibodies that belong to the first category include broadly neutralizing antibodies, such as PGT122, DH270.6 and 10-1074 (16–18).

In the closed, neutralization-resistant conformation of the HIV-1 Env, the binding site for 19b is occluded, and only becomes available upon CD-induced Env opening (19). Once CD4 has bound Env, however, the proximity to the host membrane would render the access of antibodies sterically unfavorable. As a result, despite their strong binding, 19b-like antibodies are non-neutralizing or are only able to neutralize isolates that readily sample the open conformation even in the absence of receptor binding. Despite their lack of binding to the pre-receptor bound closed Env, 19b-like antibodies are found in most individuals with HIV-1, with their elicitation likely being through the exposure of partially open Env, viral debris or shed gp120 to the immune system. Due to the conformation-selective nature of their binding, 19b-like antibodies likely play a role in driving intra-host evolution of the virus to adopt more neutralization resistant, closed conformations.

Stabilizing the closed Env conformation that only binds bnAbs and not non-neutralizing antibodies like 19b has been a longstanding goal of HIV-1 vaccine development with several strategies developed to engineer a closed Env (20–22). 19b has been extensively used in immunoassays to detect the presence of open Env conformations (19, 21–24). While the general location of the 19b epitope was known, this study now provides high resolution structures that confirm a cradle-binding mode for 19b (**Figure S2**), defining the 19b epitope at atomic level resolution and elucidating the structural basis for its broad reactivity with diverse HIV-1 isolates. Combining the structural observations with the SPR assays revealed differences in kinetics and affinity of diverse V3 peptide engagement with 19b even though the final bound poses of the V3 peptides were almost identical, providing a glimpse into the accommodation of V3 sequence and conformational diversity by 19b.

Our studies provide the structural basis for previously published reports that demonstrated the importance of certain V3 residues in 19b binding. A key role was assigned in previous studies for V3 residue 316. We found in our structures that the aromatic side chain of Phe316 was inserted at the mouth of a cavity lined by residues from the heavy and light chains. This “Phe316” cavity is reminiscent of the “Phe43” cavity that engages residue Phe43 of CD4 at the junction of the inner and outer domains of the HIV-1 Env gp120 subunit (25). Like the gp120 “Phe43” cavity, the “Phe316” cavity of 19b is malleable and can accommodate other hydrophobic/aromatic side chains in place of a Phenylalanine. Both cavities also harbor polarity through internal water molecules that can accommodate polar side chain moieties within the cavity. Thus, alongside the gp120 “Phe43” cavity, the 19b “Phe316” cavity is another example of how a single phenylalanine residue can play a dominant role in ligand binding through insertion of its aromatic side chain into an interfacial cavity to glue together different domains or subunits of the protein that they are binding to.

Through combined insights from high resolution experimental structural determination and protein structure prediction, we identified water molecules in the structure of the 19b Fab that could be critical for maintaining the conformation of the CDR L2 region and hence likely to be important for function. In the absence of these waters, AlphaFold3 failed to predict the structure of the CDR L2 region correctly. Thus, our studies provide mechanistic insights into the role of structural waters and shine a spotlight on a blind spot of modern protein prediction algorithms that don’t consider waters in their predictions. Close integration of these powerful protein structure prediction algorithms with methods that account for the role of structural waters would be vital to improve the accuracy of prediction of novel water-supported structures.

In summary, we have determined several high-resolution crystal structures of the human monoclonal antibody 19b bound to HIV-1 Env V3 peptide. Our structures have provided atomic level information on the antibody-peptide contacts including visualization of interfacial water molecules and have revealed the structural basis for the recognition of V3 peptides from diverse isolates by 19b.

## Acknowledgements

Use of the NYX beamline 19-ID at the National Synchrotron Light Source II was supported by the New York Structural Biology Center. The Advanced Light Source is a Department of Energy Office of Science User Facility under Contract No. DE-AC02-05CH11231. Thanks to Baptiste Aussedat at Chemitope Glycopeptide for V3 peptide synthesis. Thanks to Greg Buhrman for helpful discussions. This work was supported by funding from NIH R01 AI145687, R01 AI165147, U54 AI170752 and U01 AI169587.

## Author contributions

S.K.F. purified proteins, performed crystallization experiments, determined structures, performed SPR and DSF experiments; S.K.F., A.M. and P.A. performed structural analysis; N.I.N. and C-H.C. assisted with x-ray data collection, J.L. assisted with x-ray data collection, structure determination and SPR measurements; S.K.F. and P.A. prepared figures and wrote the manuscript, which all authors reviewed and approved.

## Declaration of interest

The authors declare no competing interest.

## METHOD DETAILS

### 19b heavy and light chain sequences

#### 19b light chain

##### Nucleotide sequence

gctagcaccatggagacagacacactcctgctatgggtactgctgctctgggttccaggttccactggtgacgacattgtgctgacccagtctccaccctccctgtctgcatctgtcggagacagagtcaccatcacttgccaggcgagtcaggacattagcgaccatttaagttggtttcagcagaaaccagggaaagcc cctaaactccttgtgtacggtgtatccagcttggaagcaggggtcccatctcggttcagtgtaagtggatctgggacacattttactttcaccgtcaacggcctgcagcctgaagatcttgcaacttatttctgtcagcagtatgatgatctcccgtggacgttcggcccagggaccgtggtggaagtcaaacgaactgtggctgcaccatctgtcttcatcttcccgccatctgatgagcagttgaaatctggaactgcctctgttgtgtgcctgctgaataacttctatcccagagaggccaaagtacagtggaaggtggataacgccctccaatcgggtaactcccaggagagtgtcacagagcaggacagcaaggacagcacctacagcctcagcagcac cctgacgctgagcaaagcagactacgagaaacacaaagtctacgcctgcgaagtcacccatcagggcctgagctcgcccgtcacaaagagcttcaacaggggagagtgttaggcggccgc

##### Amino acid sequence

metdtlllwvlllwvpgstgddivltqsppslsasvgdrvtitcqasqdisdhlswfqqkpgkapkllvygvssleagvpsrfsvsgsgthftftvnglq pedlatyfcqqyddlpwtfgpgtvvevkrtvaapsvfifppsdeqlksgtasvvcllnnfypreakvqwkvdnalqsgnsqesvteqdskdstysls stltlskadyekhkvyacevthqglsspvtksfnrgec*

#### 19b heavy chain

##### Nucleotide sequence

gctagcaccatggagacagacacactcctgctatgggtactgctgctctgggttccaggttccactggtgacgaggtgcagctggtacagtctgggggagccttgatccagccgggggggtccctgagactctcctgtgcagcctctggattcacttttgtcgattatgccatgagttgggtccgccaggctccagggaaggggctgcagtgggtctcaactattattggcagtggtgctgacacatactacacagactccgtgaagggccgcttcaccatctccagagacaattccaataatactgtgcatctgcaaatgaacagcctgcgagccgacgacacggccctctattattgtgtcagaggcgtctttgacctctggggccagggaacgctggtcaccgtctcctcagcctccaccaagggcccatcggtcttccccctggcaccctcctccaagagcacctctgggggcacagcggccctgggctgcctggtcaaggactacttccccgaaccggtgacggtgtcgtggaactcaggcgccctgaccagcggcgtgcacaccttcccggctgtcctacagtcctcaggactctactccctcagcagcgtggtgaccgtgccctccagcagcttgggcacccagacctacatctgcaacgtgaatcacaagcccagcaacaccaaggtggacaagagagttgagcccaaatcttgtgacaaaactcacacatgcccaccgtgcccagcacctgaactcctggggggaccgtcagtcttcctcttccccccaaaacccaaggacaccctcatgatctcccggacccctgaggtcacatgcgtggtggtggacgtgagccacgaagaccctgaggtcaagttcaactggtacgtggacggcgtggaggtgcataatgccaagacaaagccgcgggaggagcagtacaacagcacgtaccgtgtggtcagcgtcctcaccgtcctgcaccaggactggctgaatggcaaggagtacaagtgcaaggtctccaacaaagccctcccagcccccatcgagaaaaccatctccaaagccaaagggcag ccccgagaaccacaggtgtacaccctgcccccatcccgggaggagatgaccaagaaccaggtcagcctgacctgcctggtcaaaggcttctatcccagcgacatcgccgtggagtgggagagcaatgggcagccggagaacaactacaagaccacgcctcccgtgctggactccgacggctccttcttcctctata gcaagctcaccgtggacaagagcaggtggcagcaggggaacgtcttctcatgctccgtgatgcatgaggctctgcacaaccactacacgcagaagagcctctccctgtctccgggtaaatgagcggccgc

##### Amino acid sequence

metdtlllwvlllwvpgstgdevqlvqsggaliqpggslrlscaasgftfvdyamswvrqapgkglqwvstiigsgadtyytdsvkgrftisrdnsnn tvhlqmnslraddtalyycvrgvfdlwgqgtlvtvssastkgpsvfplapsskstsggtaalgclvkdyfpepvtvswnsgaltsgvhtfpavlqssgl yslssvvtvpssslgtqtyicnvnhkpsntkvdkrvepkscdkthtcppcpapellggpsvflfppkpkdtlmisrtpevtcvvvdvshedpevkfn wyvdgvevhnaktkpreeqynstyrvvsvltvlhqdwlngkeykckvsnkalpapiektiskakgqprepqvytlppsreemtknqvsltclvk gfypsdiavewesngqpennykttppvldsdgsfflyskltvdksrwqqgnvfscsvmhealhnhytqkslslspgk*

### 19b expression and purification

Plasmids encoding the heavy and light chains of IgG 19b were mixed in a 1:1 molar ratio for transient transfection in Expi293i cells at 2.5 x10^6 cells/mL followed by addition ExpiFectimine 18 hours later. Supernatant was harvested after 5 days and filtered at 0.22 μm. Supernatant was poured over Protein A resin using gravity chromatography. 19b was eluted using IgG Elution buffer (ThermoScientific) having adjusted to pH 7 using Tris pH 8. Antibody was concentrated to 35 – 73 mg/mL and stored – 80 C before Fab digest. LysC (Roche) was mixed in 1:2000 mass ratio and reaction occurred at 37 °C for 4 hours. 1x protease inhibitor solution (Pierce) was added to stop reaction and mixture stored at 4 °C. A Protein A Plus Agarose NAb spin column was used to purify 19b Fab. Eluted Fab was concentrated and further purified by size-exclusion chromatography (Superose 6 Increase, 10/300 GL column) in 1x PBS pH 7.4, peak fractions were pooled and 19b Fab was concentrated to approx. 50 mg/mL. Aliquots were snap cooled in liquid N2 and stored at – 80 °C until use.

### SPR (Surface Plasmon Resonance)

SPR binding measurements were taken at room temperature using a Cytiva Biacore T-200 instrument. V3 peptides with biotin moiety at the C-terminus were diluted to 10 nM in HBS/3 mM EDTA/0.05 % P-20 and flowed over a high-affinity streptavidin (SA) sensor chip for attachment with final response units (RUs) in the range of 50 - 200. 19b Fab was injected using the single cycle kinetics mode with 5 concentrations per cycle in the range of: 20 nM, 10 nM, 5 nM, 2.5 nM, 1.25 nM, 0.625 nM and 0.3125 nM. A contact time of 60s and a dissociation time of 60 s at a flow rate of 50 mL/min was used. For 5768-p27 peptides, a dissociation time of 2 h was needed. The surface was regenerated after the last injection with 3 pulses of both a 50 mM NaOH + 1M NaCl solution and for 5768(x)-p27 peptides, supplemental 50 %(v/v) ethylene glycol solution for 10 s at 100 mL/min. Flow cell 1 was used as reference. Blank sensorgrams were obtained by injection of the same volume of HBSEP + buffer in place of IgG solutions. Sensorgrams were reference subtracted and corrected with corresponding blank curves. Sensogram data were analyzed using the BiaEvaluation software (Cytiva).

### DSF (differential scanning fluorimetry)

Samples were diluted to 1.25 mg/mL and run in triplicate on the Prometheus Panta (Nanotemper). Ramp was run at 1 °C / min, from 25 °C to 95 °C with excitation at 280 nm and emission detected at 330 nm and 350 nm. PR.Panta Control v1.6.3 and PR.Panta Analysis v1.6.3 were used to collect and analyze the samples respectively.

### X-ray crystallography

19b Fab and V3 peptides were mixed in a 1:1.3 molar ratio as follows: for JR-FL, 92BR020 and 5768(x)-p27, 1.0 mg of dried powder of peptide was weighed then dissolved in 50 mg/mL of 19b Fab in 1x PBS solution. Complexes were incubated 1.5 – 3 hours at 4 °C. Unbound 19b Fab was defrosted for 1.5 h at 4 °C before crystallization trials. Oryx4 (Douglass Instruments) and Mosquito (SPT Labtech) liquid handlers were used for the sitting drop vapor diffusion method. An Integra 300 was used to dispense 70 uL of the screening solutions to the plates (Art-Robbins Intelliplate 3 well). 3 drops were used for each condition: 0.2 μL protein + 0.2 μL condition, 0.15 μL protein + 0.25 μL condition and 0.25 μL protein + 0.15 μL condition. Crystal Screen Cryo and PEG/Ion screens from Hampton were used to screen for initial crystals. Crystal growth was monitored by taking drop images on a Jansi UVEX robot imager in bright field and UV. Crystals were harvested and cryoprotectant consisted of the condition supplemented with 30 %(v/v) glycerol.

X-ray diffraction data collection and indexing details are found in **Table S1**. Aimless was employed to create the .mtz file used for PHASER/MR in PHENIX, followed by PHENIX.refine. To create the 19b Fab structure, 17b Fab (PDB 7KLC) (26) was used as the initial Fab model. Following which, the sequence of 19b was validated by GeneImmune (Supplement Figure 2). For initial model building, a single pdb file was created separating the variable region of the 17b light and heavy chains using PDB tools in PHENIX. The same was done for the constant region of the light and heavy chains of 17b. The V3 peptide model was built into the difference density initially for the 19b Fab - JR-FL complex, then modified for the other peptide structures. Omit maps were generated for the V3 peptides in complex with 19b. COOT v0.9.8.91,xf Pymol v2.2.3, PHENIX 1.20.1-4487 were used for model building, refinement and figure making (27–29).

---

**Figure S1.**
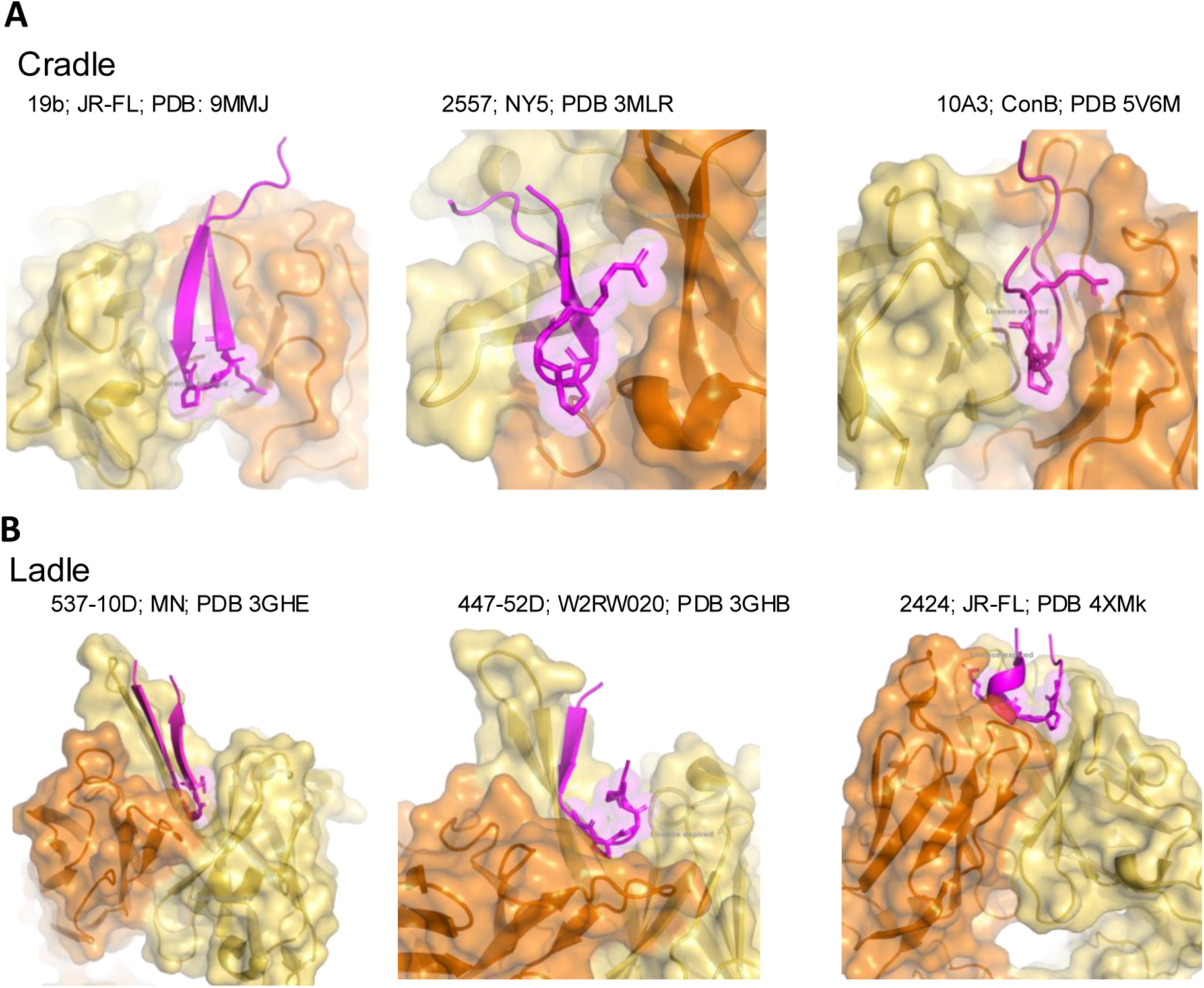
19b adopts the cradle binding mode. Representative examples of V3 antibodies that adopt the (A) cradle and (B) ladle binding mode.

**Figure S2.**
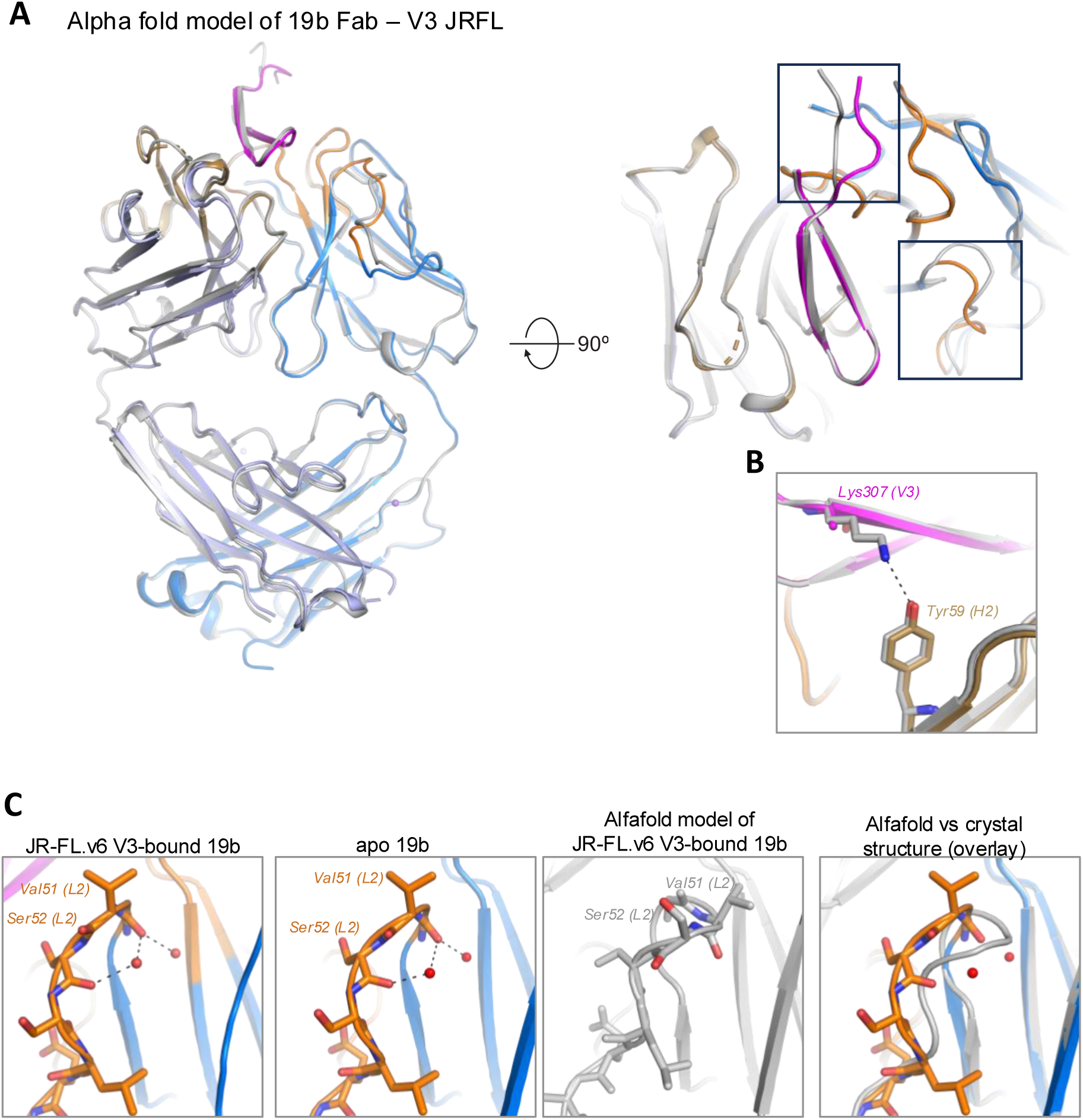
AlphaFold prediction of 19b crystal structures. (A) Superposition of 19b Fab – V3 JR-FL crystal structure with AlphaFold predicted model has Cα RMSD of 0.477 Å. AlphaFold model (grey) was predicted before deposit of the 19b crystal structures into the Protein Data Bank. (B) Zoomed in view of the V3 Lys307 – 19b Tyr59 (H2) hydrogen bond predicted. Side chain for Lys307 was not modeled in the crystal structure due to lack of side chain electron density. (C) Zoomed in views of the 19b CDR L2 loop for the Left to right. JR-FL.v6 V3-bound crystal structure, crystal structure of apo 19b Fab, AlphaFold predicted structure of JR-FL.v6 V3-bound 19b Fab complex, and overlay of the AlphaFold predicted and experimentally determined structures of the JR-FL.v6 V3-bound 19b Fab complex.

**Table S1.**
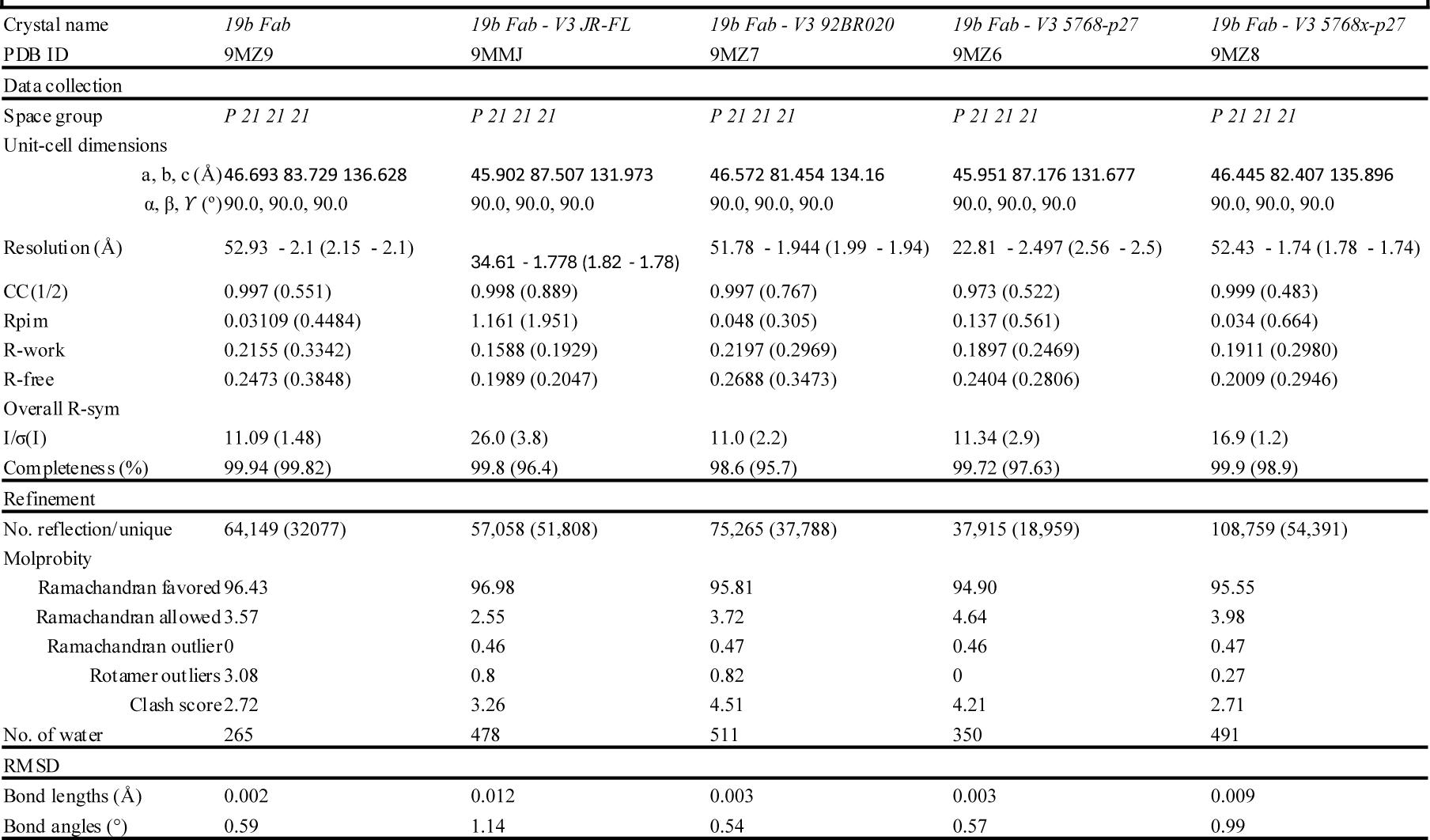
Crystal lograph ic data collection and refinement.

**Table S2:**
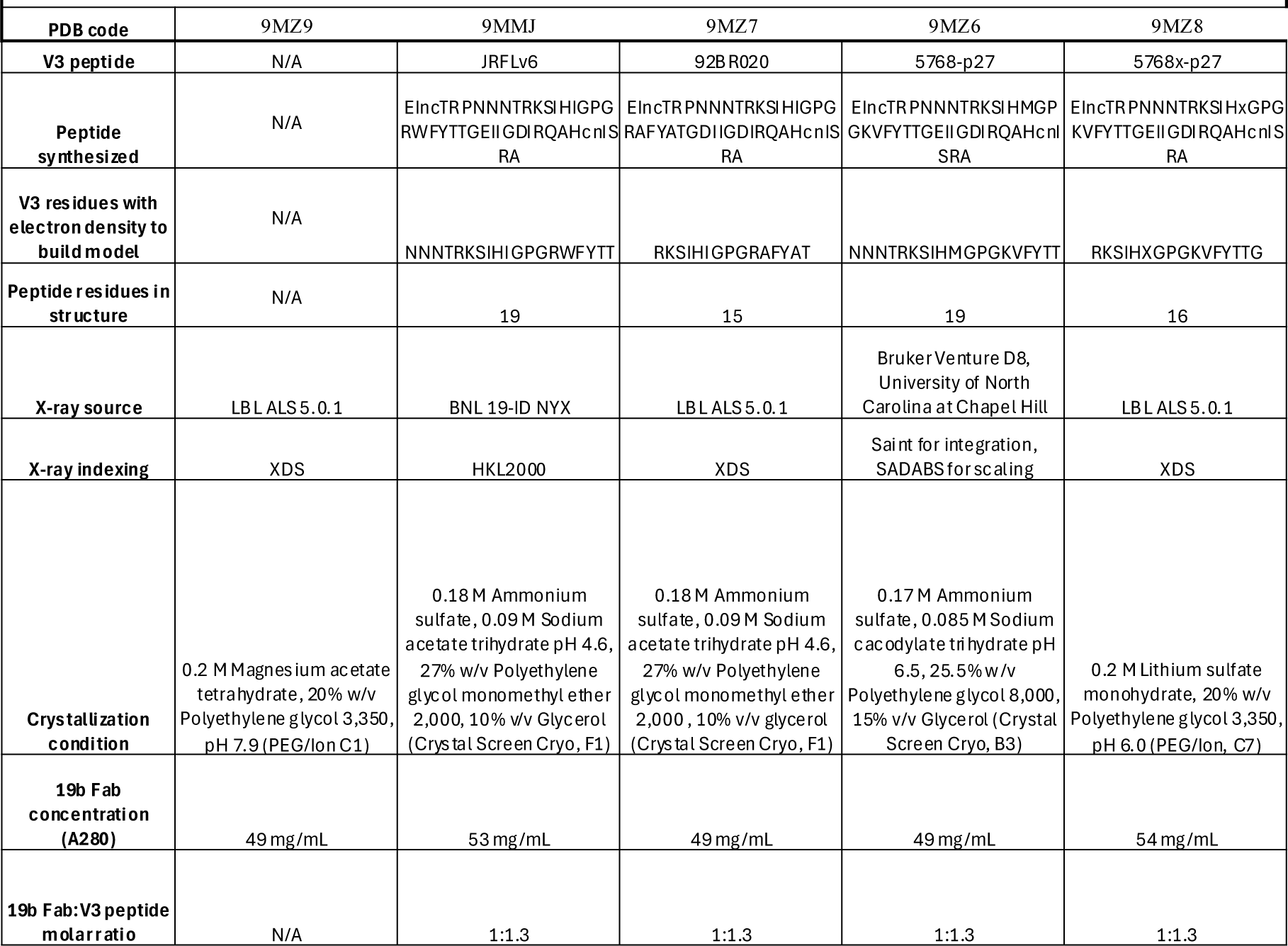
19b Fab crystallization conditions.

**Table S3:** RMSDs of 19b Fab strcutres compared to 19b Fab - V3 JRFL.

| Table S3: RMSDs of 19b Fab strcutres compared to 19b Fab - V3 JRFL |  |  |  |  |  |  |  |  |  |  |
| --- | --- | --- | --- | --- | --- | --- | --- | --- | --- | --- |
| Peptide | PDB Code | Light Chain | Heavy Chain | CDR L1 | CDR L2 | CDR L3 | CDR H1 | CDR H2 | CDR H3 | V3 |
| Ligand-free | 9MZ9 | 0.453 | 1.039 | 0.094 | 0.143 | 0.117 | 0.069 | 0.121 | 0.295 | N/A |
| 92BR020 | 9MZ7 | 0.478 | 0.517 | 0.087 | 0.143 | 0.075 | 0.119 | 0.104 | 0.104 | 0.208 |
| 5768-p27 | 9MZ6 | 0.144 | 0.147 | 0.097 | 0.113 | 0.079 | 0.106 | 0.083 | 0.111 | 0.201 |
| 5768x-p27 | 9MZ8 | 0.44 | 0.983 | 0.087 | 0.189 | 0.041 | 0.091 | 0.062 | 0.066 | 0.202 |

## Notes

### Competing Interest Statement

The authors have declared no competing interest.

### Summary of Updates

Corrections made to Figure 5 legend. Corresponding figure call outs updated in the text. Minor typos corrected.

